# Cells in the Polyaneuploid Cancer Cell (PACC) state have increased metastatic potential

**DOI:** 10.1101/2022.09.16.508155

**Authors:** Mikaela M. Mallin, Nicholas Kim, Mohammad Ikbal Choudhury, Se Jong Lee, Steven S. An, Sean X. Sun, Konstantinos Konstantopoulos, Kenneth J. Pienta, Sarah R. Amend

**Affiliations:** Cellular and Molecular Medicine Graduate Training Program, Johns Hopkins School of Medicine, Baltimore, MD, USA; Cancer Ecology Center, James Buchanan Brady Urological Institute, Johns Hopkins Medical Institute, Baltimore, MD, USA; Rutgers Institute for Translational Medicine and Science, New Brunswick, NJ, USA; Department of Mechanical Engineering, Johns Hopkins University, Baltimore, MD, USA; Department of Chemical and Biomolecular Engineering, Johns Hopkins University, Baltimore, MD, USA

**Keywords:** Polyaneuploid Cancer Cells (PACCs), Polyploid Giant Cancer Cells (PGCCs), Chemotaxis, Deformability, Vimentin

## Abstract

Although metastasis is the leading cause of cancer deaths, it is quite rare at the cellular level. Only a rare subset of cancer cells (∼1 in 1.5 billion) can complete the entire metastatic cascade: invasion, intravasation, survival in the circulation, extravasation, and colonization (i.e. are metastasis competent). We propose that cells engaging a Polyaneuploid Cancer Cell (PACC) phenotype are metastasis competent. PACCs are enlarged, non-dividing cells with increased genomic content that form in response to stress. Single-cell tracking using time-lapse microscopy reveals that PACCs are more motile than nonPACCs. Additionally, PACCs exhibit increased capacity for environment-sensing and directional migration in chemotactic environments, predicting successful invasion. Magnetic Twisting Cytometry and Atomic Force Microscopy reveal that cells in the PACC state display hyper-elastic properties like increased peripheral deformability and maintained peri-nuclear cortical integrity that predict successful intravasation and extravasation. Furthermore, four orthogonal methods reveal that PACCs have increased expression of Vimentin, a known hyper-elastic biomolecule. Lastly, anoikis-resistance assays and detection of PACCs in the blood of a patient with metastatic castrate-resistant prostate cancer using a selection-free circulating tumor cell detection platform reveal that PACCs are capable of surviving in the circulation. Taken together with the knowledge that PACCs are capable of eventual depolyploidization and progeny formation (as a potential route to colonization), these data support PACCs as candidate metastasis-competent cells worthy of further analysis.

## Introduction

Metastasis describes the development of secondary malignant growths at a distant site and is responsible for 90% of cancer related deaths [1].The metastatic cascade describes the steps that cancer cells take as they travel through the body: invasion, intravasation, survival in the circulation, extravasation, and colonization. To form lethal metastatic lesions, cancer cells in the primary tumor first must acquire an invasive capacity to penetrate surrounding extracellular matrices. Second, the cells must enter the circulatory system via a process called intravasation, extending into neighboring vascular tissues and maneuvering between endothelial cells. This requires altered cancer cell cytoskeletal dynamics that increase deformability yet simultaneously preserve nuclear envelope integrity. Once in the circulatory system, the cells, now called a circulating tumor cells (CTCs), must evade anoikis, a form of programmed cell death initiated upon detachment from extracellular matrix. Fourth, the cells must extravasate out of the circulatory system into distant secondary organs. Lastly, the cells, now called disseminated tumor cells (DTCs), must survive and subsequently proliferate enough to form an overt metastatic lesion [2]. A cancer cell must be capable of successfully completing all five steps of the metastatic cascade to create a metastatic tumor. Cells with this capability are termed metastasis-competent cells.

Although metastasis is common at the organismal level (roughly 30% of cancer patients will develop metastases) [1], clinically-based mathematical modeling predicts that at the cellular level, metastasis is actually an extremely rare event. Estimates based on the number of CTCs detectable in a patient’s blood, the number of DTCs detected in their bone marrow (a common metastatic site for multiple cancers), and the number of clinically detectable metastatic lesions suggest that only 1 of every 1.5 billion CTCs successfully completes the metastatic cascade [3]. Note that these calculations exclude cancer cells that never acquire motility, meaning that the proportion of metastasis-competent cells in the primary tumor is even lower. The identity of the rare subpopulation of cancer cells with true metastatic competency remains an open research question.

Conventionally, it is understood that cellular stress results in activation of a canonical stress response, resulting in a brief pause in cell cycle (during which damage is repaired) followed by either swift cell cycle reactivation or apoptosis [4, 5]. The Polyaneuploid Cancer Cell (PACC) state describes a recently appreciated third stress-response fate [6-15]. The PACC state is a transient, adaptive state adopted by some cells in response to stress [16]. The PACC state is characterized by physically enlarged cell size, increased genomic content, and lack of cell division. Polyaneuploid specifically describes the stress-induced polyploidization of an already aneuploid cancer cell genome. Cells in the PACC state (PACCs) have been described using other names, including Polyploid Giant Cancer Cells, Multinucleated Giant Cancer Cells, and Pleomorphic Cancer Cells, among others [17-20]. Regardless of nomenclature, these types of cells have repeatedly been implicated in metastasis and chemotherapeutic resistance [21-39].

PACCs arise in response to many stressors, including hypoxia, acidity, ionizing radiation, and various classes of chemotherapeutic drugs, including cisplatin, docetaxel, and etoposide [6, 40]. Moreover, the PACC state has been described in multiple cancer types, including prostate cancer lines PC3, DU145, and LNCaP, breast cancer line MDA-MB-231, and ovarian cancer cell lines HEY and SKOv3, among many others [25]. PACCs can be found at low numbers in untreated cultures, likely reflecting low but ever-present cellular stress inherent to routine cell culture practices. Application of additional tumor microenvironmental stressors causes a marked increase in the number of PACCs present. While some of the cells exposed to the applied stress engage in canonical stress-response programs, a subset of cells exposed to the applied stress initiate a Polyaneuploid Transition (PAT) to access the PACC state. Upon accession of the PACC state, PACCs retain high levels of cellular functionality, including operative respiration, biosynthesis, digestion, absorption, secretion, homeostasis, transport, and movement [41]. Although cells in the PACC state are nonproliferative, they increase markedly in size at a steady rate. Eventually, PACCs undergo depolyploidization (i.e. re-engage in reductive cell division) and exit the PACC state by producing proliferative nonPACC cancer cell progeny of normal physical size and typical genomic content [40, 42-50].

Cells in the PACC state have been identified histologically in various human cancer types in multiple steps of the metastatic cascade. Presence of PACCs within primary tumors is generally associated with poorer prognosis [51, 52]. Recently, it was shown that existence of PACCs in the primary tumors of men with prostate cancer who underwent radical prostatectomy with curative intent is predictive of lower metastasis-free survival [53]. Additionally, rapid autopsy of 5 men with metastatic prostate cancer revealed presence of PACCs in all distant metastatic lesions analyzed [54]. In animal models, highly metastatic prostate cancer cells selected by serial metastatic passage in mice are highly enriched for PACCs [55]. Taken together, these data suggests that cells in the PACC state may be metastatically competent. This hypothesis predicts that PACCs can invade, intravasate, and survive in the circulation. To test these predictions, we quantified the motility, deformability, and anoikis resistance of cisplatin-induced PACCs derived from the PC3 prostate cancer cell line.

## Results

### Cancer cells undergo a Polyaneuploid Transition (PAT) in response to chemotherapeutic stress

The PC3 prostate cancer cell line was treated with 6 uM of cisplatin for 72 hours, after which cisplatin-containing media was replaced with fresh media. After treatment, all cells had characteristics of PACC morphology (increased cell size, increased nuclear size, distinctive nuclear morphology, and increased peri-nuclear granularity including increased lipid droplet content), indicating they had initiated PAT (Figure 1a). Polyploidization was validated by flow cytometry on a subset of cells to measure the relative DNA content of treated cells compared to nontreated controls. The cell cycle plots of treated cells demonstrate a clear rightward shift compared to the typical cell cycle plot of nontreated controls, indicated a >G2 ploidy (Figure 1b). All remaining treated cells were maintained in culture and monitored for an additional 10 days via in-incubator time-lapse microscopy. Treated cells did not undergo any cellular division during this time, but they did continue to increase in size (i.e. 2D surface area) throughout the duration of the experiment. Treated cultures continued to show evidence of cell death until day 10 after treatment release, after which all surviving cells are definitively in the PACC state (Supplemental Figure 1). All following experiments were completed with PACCs maintained in culture without dividing for 10 days post cisplatin-treatment, unless otherwise indicated.

**Fig. 1.**
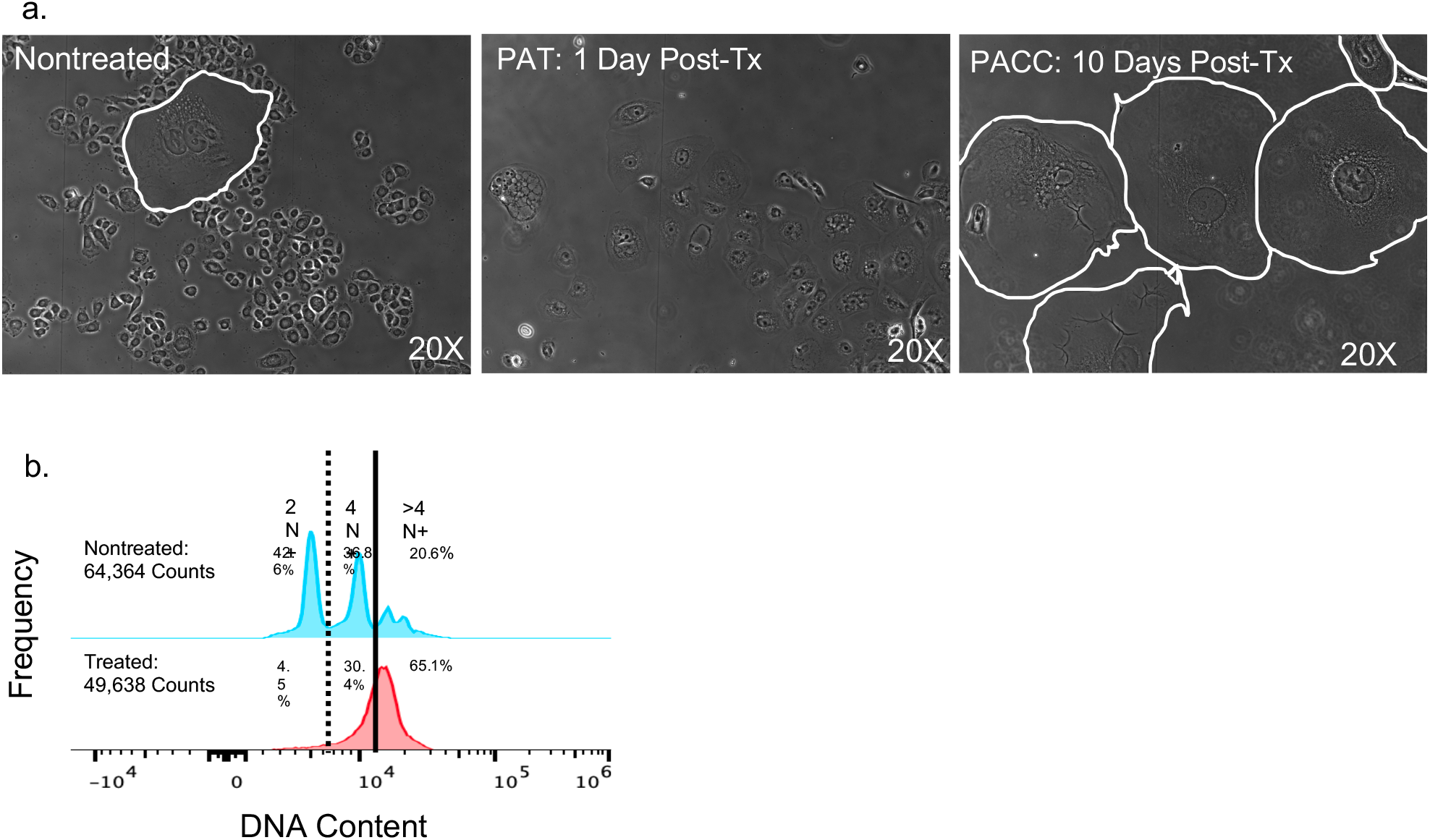
Cancer cells undergo a Polyaneuploid Transition in response to applied stress: a) 2D morphology of PACC induction in PC3 cells using a 72 hour dose of 6 uM cisplatin. Cell populations are untreated, undergoing a PAT 1 day post-treatment, or have entered a definitive PACC state 10 days post-treatment. Acquired by 20X phase microscopy. PACC cell borders are demarcated in white. b) Relative DNA content of nontreated cells vs. treated cells 1 day post-treatment. Acquired by flow cytometry

### Cells in the PACC state have increased motility

Motility is critical for metastasis initiation and is prerequisite to the first step of the metastatic cascade. To directly quantify the motility of cells in the PACC state compared to nonPACC parental cells, we performed time lapse microscopy followed by single cell tracking using ImageJ. Motility videos of nonPACCs and PACCs are included as Online Resource 1 and 2, respectively. All cells were exposed to uniform 20% FBS-supplemented media conditions, denoted as +/+.

While cells in the PACC state travelled lesser accumulated distances (p value = 0.0002) than nonPACC parental cells (Figure 2a), Cells in the PACC state travelled much greater Euclidean distances (p value = 0.0002) than nonPACC parental cells (Figure 2b). Accumulated distance describes the total distance traveled by a cell, and Euclidean describes the net distance traveled by a cell (i.e. the length of the line connecting the starting and ending points). Spider plots tracking the (X,Y) coordinates of each cell’s trajectory mapped on a Cartesian plane wherein all cell locations at Time 0 are translated to the origin (0,0) reveal that the observed increase in accumulated distance travelled by nonPACC cells is almost entirely non-migratory: the cells’ motility is mostly in place -- no large net distances are travelled (Figure 2c). Thus, Euclidean distance is a more metastasis-relevant measure of invasive potential than accumulated distance. The ratio of Euclidean distance over accumulated distance can be used to calculate directness of movement. A directness metric of 1 describes movement in a perfectly direct (i.e. straight) line, while a directness metric approaching 0 indicates totally indirect cell movement. In our analyses, PACCs moved more directly (p value < 0.0001) than nonPACC parental cells (Figure 2d).

**Fig. 2.**
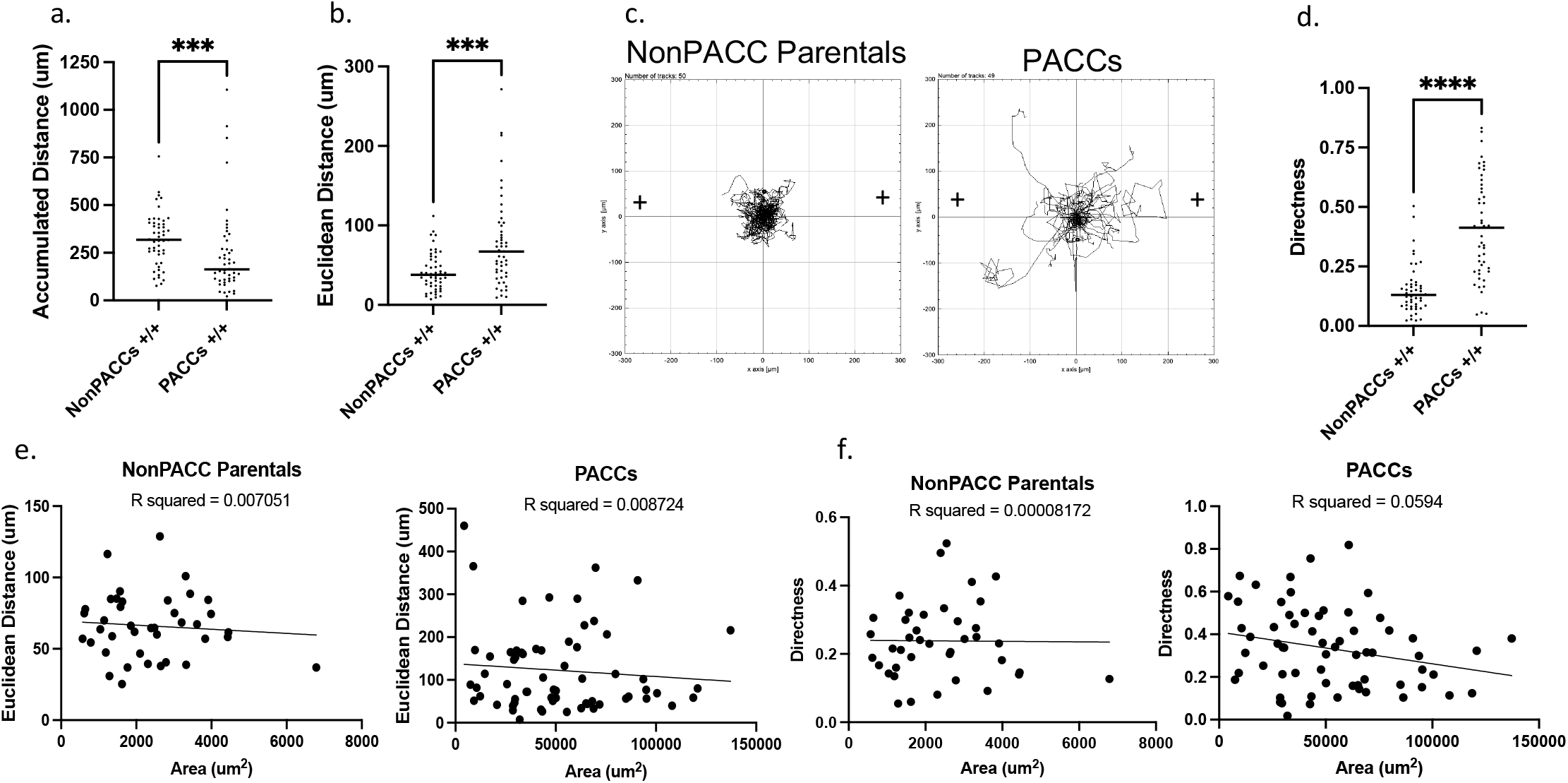
Cells in the PACC state are more motile: a-b) Quantification of the A) accumulated distance and B) Euclidean distance travelled by nonPACC parental cells or PACCs in uniform 20% FBS-supplemented media conditions (+/+) throughout a 24 hour time lapse. c) Spider plots depicting the motility tracks of single nonPACC parental cells (n=50) or PACCs (n=49) in uniform 20% FBS-supplemented media conditions (+/+) throughout a 24 hour time lapse. d) Directness of travel demonstrated by nonPACC parental cells or PACCs in uniform 20% FBS- supplemented media conditions (+/+) throughout a 24 hour time lapse. e) Linear regression comparing the Euclidean distance travelled and 2D cell surface area of nonPACC parental cells and PACCs. f) Linear regression comparing the Directness and 2D cell surface area of nonPACC parental cells and PACCs.

To test if increased PACC motility is attributable to the increased cell size of the PACC population, we performed a linear regression comparing the 2D surface area of each cell to its specific motility metrics. No correlation between cell size and cell motility within nonPACC parental cell populations or within PACC populations was found (Figure 2e). All linear regressions returned nonsignificant R^2^ values when modeling both the relationship between cellular area and Euclidean distance for both nonPACC parental cells (R squared = 0.007051) and PACCs (R squared = 0.008724) as well as the relationship between cellular area and directness for both nonPACC parental cells (R squared = 0.00008172) and PACCs (R squared = 0.0594) (Figure 2e and 2f).

### Cells in the PACC state demonstrate a directional response to a chemotactic FBS gradient

A motile cell’s speed (ergo, distance travelled) and directness are considered its kineses, a term borrowed from foraging and movement ecology used to describe basic movement parameters. Cellular movement is often kineses-directed, meaning changes in a cell’s location are directly due to cell-intrinsic adjustments to its speed and/or direction, rather than due to some extrinsic applied force. Kineses-directed movement occurs in response to local resource levels, such as directional chemotaxis up a nutrient gradient [56].

To test if PACCs are capable of resource sensing (i.e. perform kineses-directed movement, or directional chemotaxis), we performed time lapse microscopy and single cell tracking along an FBS gradient of 0-20% FBS (-/+). Positive control cells were cultured in uniform 20% FBS conditions (+/+) and negative control cells were cultured in uniform serum-free conditions (-/-). As we had previously observed, PACCs always moved greater Euclidean distances than nonPACC parental cells under all experimental conditions (+/+ p value = 0.0002, -/- p value < 0.0001, -/+ p value < 0.0001) (Figure 3a). Furthermore, cellular migrations were more direct (i.e. persistent) in PACCs than nonPACC parental cells under all experimental conditions (+/+ p value < 0.0001, -/- p value < 0.0001, -/+ p value < 0.0001) (Figure 3b). Spider plots tracking the (X,Y) coordinates of each cell’s trajectory indicate that PACCs respond more robustly to presence of an FBS gradient than nonPACC parental cells: A greater number of PACCs travel greater distances up the FBS gradient than nonPACC parental cells (Figure 3c).

**Fig. 3.**
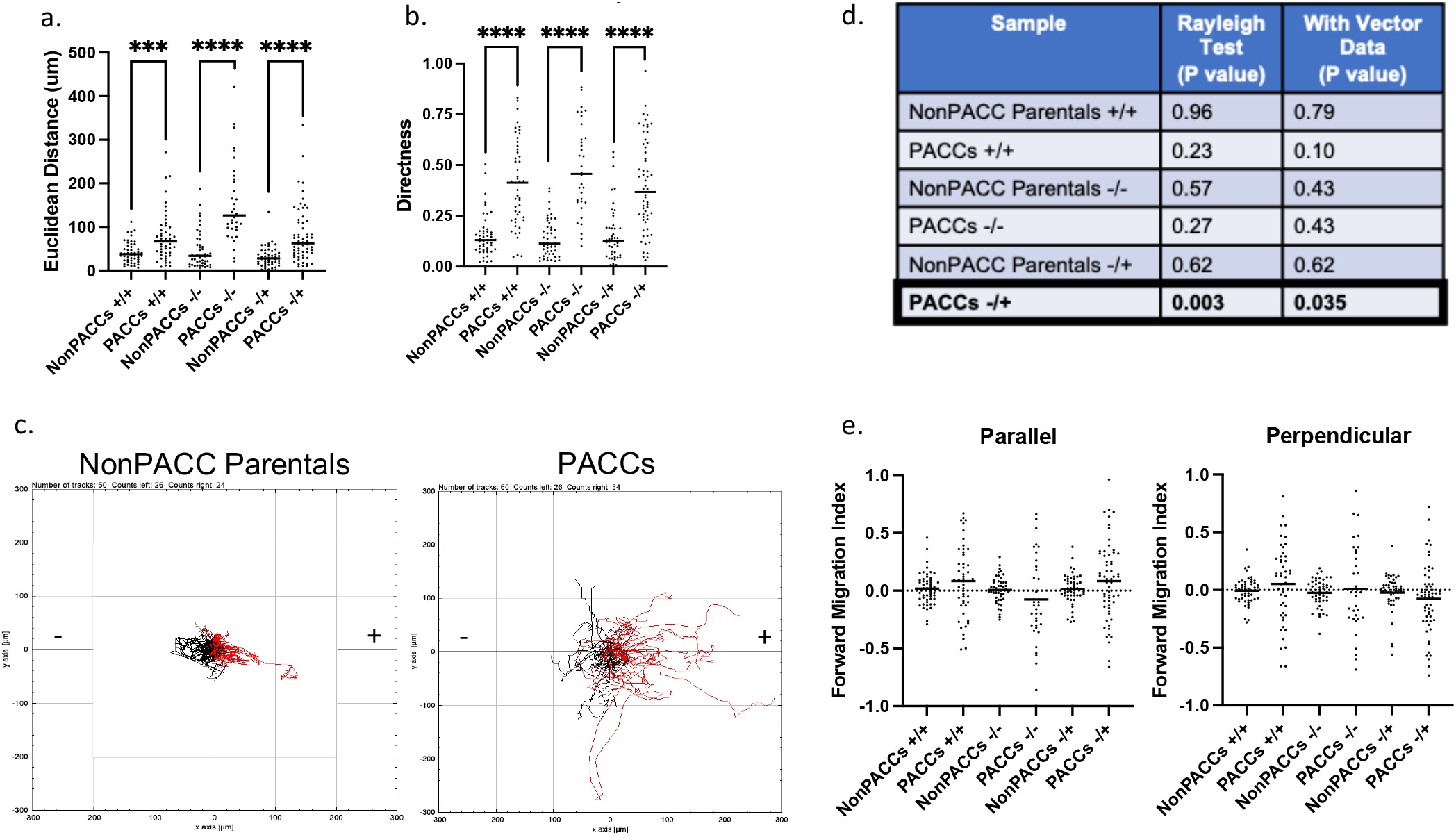
Cells in the PACC state demonstrate a directional response to a chemotactic FBS gradient: a-b) Quantification of the Euclidean distance travelled by or B) directness of movement of nonPACC parental cells or PACCs in either uniform 20% FBS-supplemented media conditions (+/+), uniform serum-free media conditions (-/-), or a chemotactic gradient of 0—20% FBS-supplemented media (-/+) throughout a 24 hour time lapse. c) Spider plots depicting the motility tracks of single nonPACC parental cells (n=50) or PACCs (n=60) in a chemotactic gradient of 0—20% FBS-supplemented media (-/+) throughout a 24 hour time lapse. Red tracks indicate cells that had a net migration toward up the FBS gradient. d) Results of the Rayleigh Test and Rayleigh Test with Vector Data for approximation of chemotactic behavior in nonPACC parental cells and PACCs. e) Parallel (to FBS gradient) and Perpendicular (to FBS gradient) Forward Migration Indices for assessment of chemotactic behavior/chemotaxis in nonPACC parental cells and PACCs

To assess if PACCs exhibited directional chemotaxis, we performed the Rayleigh Test and the Test of Forward Migration Indices. The Rayleigh Test is a statistical test for the uniformity of circularly distributed data (here, the spider plots’ final X,Y coordinate of a cell’s trajectory) [57]. The null hypothesis is circular uniformity. Final coordinates distributed equally around the origin indicate lack of directional response to the FBS gradient and will fail to reject the null, producing a nonsignificant p value. Final coordinates distributed nonequally around the origin indicate presence of a directional response and will reject the null, producing a significant p value. The Rayleigh Test for vector data also includes a parameter of distance, measured between the final coordinate and the origin. To define the movement of cells as chemotactic, the result of the Rayleigh tests for the gradient condition must be significant while the results for the control conditions must be nonsignificant. Both the Rayleigh Test and the Rayleigh Test for vector data were only significant for PACC movement under gradient conditions (Figure 3d). This indicates that PACCs exhibited directional chemotaxis in response to an FBS gradient but that nonPACC parental cells did not.

The Test of Forward Migration Indices quantifies the directional efficiency of cellular movement in a defined direction. The Parallel Forward Migration Index (|| FMI) describes the movements of cells in planes parallel to that of a present gradient. Positive || FMI values represent “forward” movement up the gradient, while negative || FMI values represent “backwards” movement down the gradient. The Perpendicular Forward Migration Index (⊥ FMI) describes the movements of cells in planes perpendicular to that of a present gradient. To define the movement of cells as chemotactic, i) the || FMI of the gradient condition must be significantly higher than the || FMI of both control conditions, ii) the || FMI of the gradient condition must be significantly higher than the ⊥ FMI of the gradient condition, which must be close to zero, and iii) the || FMI and ⊥ FMI of both control conditions must be close to zero. Our data indicate that directional chemotaxis in response to an FBS gradient is apparent in PACCs but not in nonPACC parental cells (Figure 3e).

### Cells in the PACC state have altered cytoskeletal stiffness

The second step of the metastatic cascade, intravasation, requires substantial morphological deformation as invasive cells maneuver between endothelial cell junctions. Cytoskeletal stiffness is an important cellular property that changes during migration and can be used as a proxy measure of deformability. Cytoskeletal stiffness can be described and measured in two ways: intracellular network stiffness and cortical stiffness [58].

Magnetic Twisting Cytometry (MTC) was used to measure the intracellular network stiffness of PACCs and nonPACC parental cells. Using MTC, cells in the PACC state showed decreased intracellular network stiffness compared to nonPACC parental cells. This result indicates that cells in the PACC state are more deformable than nonPACC parental cells (Figure 4a).

**Fig. 4.**
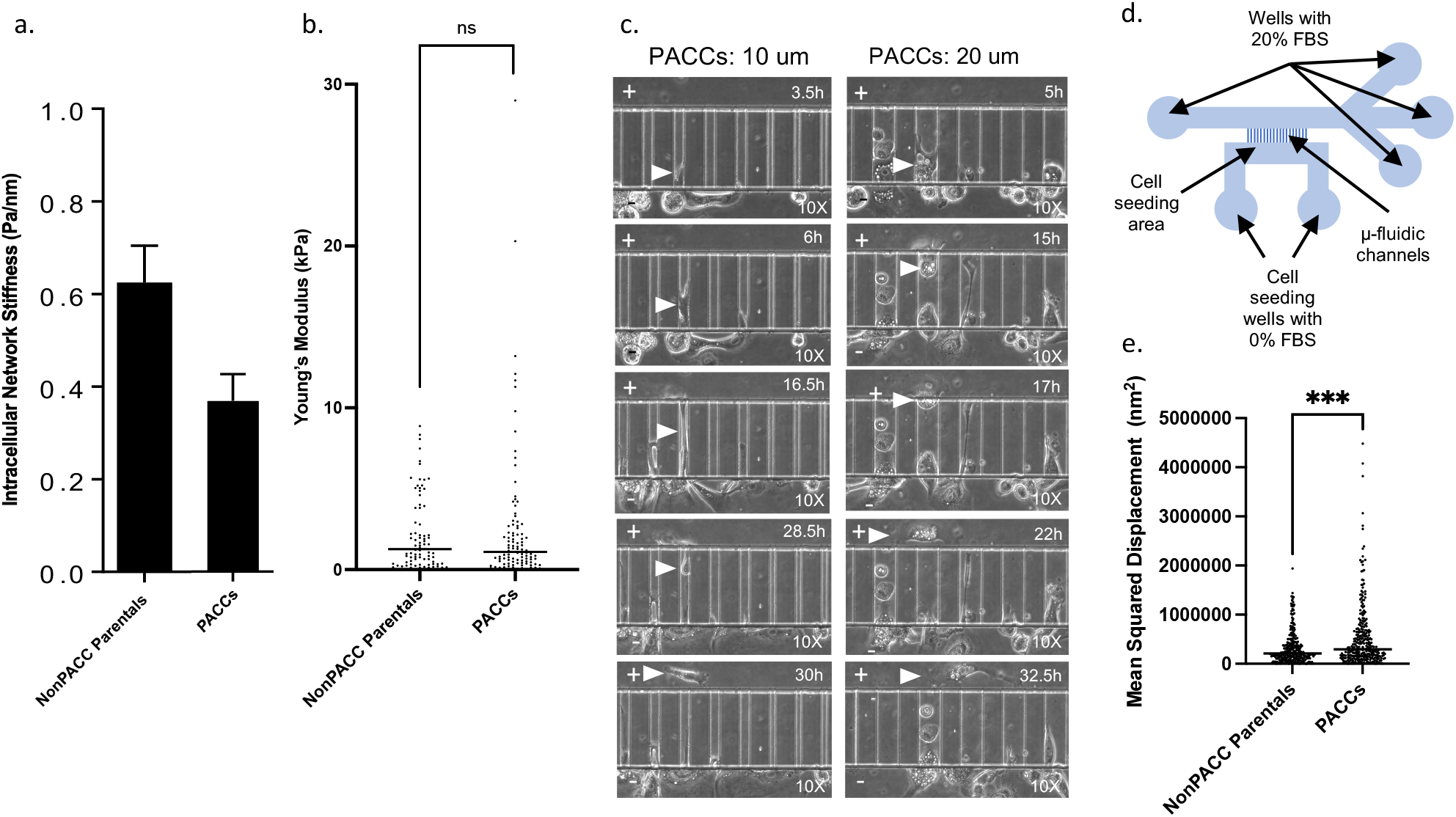
Cells in the PACC state have altered cytoskeletal stiffness and increased cytoskeletal dynamics: a) Intracellular network cytoskeletal stiffness of nonPACC parental cells PACCs acquired via MTC using ferromagnetic RGD- linked beads. b) Young’s Modulus (or cortical stiffness) of the peri-nuclear region of nonPACC parental cells and PACCs acquired using AFM. c) Demonstration of functional deformability in PACCs: Evidence of simultaneous migration and deformation of PACCs through 10 um and 20 um wide channels of a custom invasion channel microfluidic device throughout a 36 hour time-lapse. d) Schematic of the microfluidic device. e) Mean Squared Displacement of spontaneous movement of RGD-linked beads attached to the cytoskeleton of nonPACC parental cells and PACCs throughout a 5 minute time-lapse

Atomic Force Microscopy (AFM) was utilized as an orthogonal approach to further evaluate cortical stiffness in the perinuclear region of PACCs and nonPACC parental cells. Cortical stiffness is reported as Young’s modulus (E, or also elastic modulus), where a larger Young’s modulus indicates a stiffer, or less deformable, cell. By AFM there was no significant difference between the perinuclear Young’s modulus (i.e. cortical stiffness) of PACCs and that of nonPACC parental cells (p value = 0.5765) (Figure 4b).

Given the discrepancies between our MTC and AFM data, we sought to investigate the functional deformability of PACCs. To directly assess functional deformability, we simultaneously tracked cell movement and deformation through a custom-built microfluidic invasion chamber [59, 60]. In this device, cells are seeded in a primary chamber that is connected to a secondary chamber via narrow invasion channels of varying sizes (3, 6, 10, 20, and 50 um). Cells invade from the primary chamber to the secondary chamber by squeezing through the invasion channels (Figure 4d). A 0-20% FBS gradient was applied across the invasion channels to stimulate directional movement. Time lapse imaging demonstrated that cells in the PACC state are capable of functional deformability; PACCs can deform while simultaneously moving into and through the 10 um and 20 um invasion channels (Figure 4c). Videos of PACC functional deformability through 10 um and 20 um channels are included as Online Resources 3 and 4, respectively.

### Cells in the PACC state exhibit increased cytoskeletal dynamics

The cytoskeleton is the primary organelle responsible for cellular migration and deformability. It has been reported that increased measures of cytoskeletal dynamics correlate with increased cellular migration and deformability [61]. The cytoskeletal dynamics of PACCs and nonPACC parental cells were evaluated by functionalizing RGD-coated microbeads to the cells’ cytoskeleton through cell surface integrin receptors and tracking individual bead displacement via short-range time lapse microscopy. Spontaneous bead motions report the underlying remodeling dynamics of the cytoskeletal network to which the beads are attached (i.e., greater cytoskeletal rearrangement dynamics are reflected as greater RGD bead displacement, reported as mean squared displacement). Cells in the PACC state had significantly greater RGD bead displacement than nonPACC parental cells (p value = 0.0002), indicating that PACCs rearrange their cytoskeleton more dynamically (Figure 4e). This data is consistent with the direct observations of increased motility and increased deformability in PACCs.

### Cells in the PACC state have increased Vimentin expression

Vimentin (VIM) is a Type III Intermediate filament component of the cytoskeleton and a canonical marker of Epithelial-to-Mesenchymal transition (EMT). VIM has been implicated in two major cellular functions: mesenchymal cell motility and preservation of nuclear-cytoskeletal integrity, especially in response to compressive stress. It has also been found to play an important role in cell spreading on firm 2D substrates [62-70].

We assessed VIM levels at both the RNA and protein level in cisplatin-induced cells at various time points throughout the PAT as well as in cells in a definitive PACC state. mRNA NanoString analysis of treated and nontreated cells immediately following 72-hour exposure to IC50 doses of either cisplatin, docetaxel, or etoposide showed a 3.5-5 fold increase in *VIM* mRNA upon entry into the PAT (Figure 5a). This was validated by RT-qPCR analysis: cisplatin-treated cells had increased *VIM* expression compared to nontreated controls. *VIM* expression steadily increased with time post-treatment throughout the PAT and into the definitive PACC state (10 days post-treatment) (Figure 5b). At the protein level, western blot analysis of treated and nontreated cells likewise showed that VIM protein expression increased with cisplatin-induction and continued to steadily increase with time post-treatment throughout the PAT and into the definitive PACC state (10 days post-treatment) (Figures 5c).

**Fig. 5.**
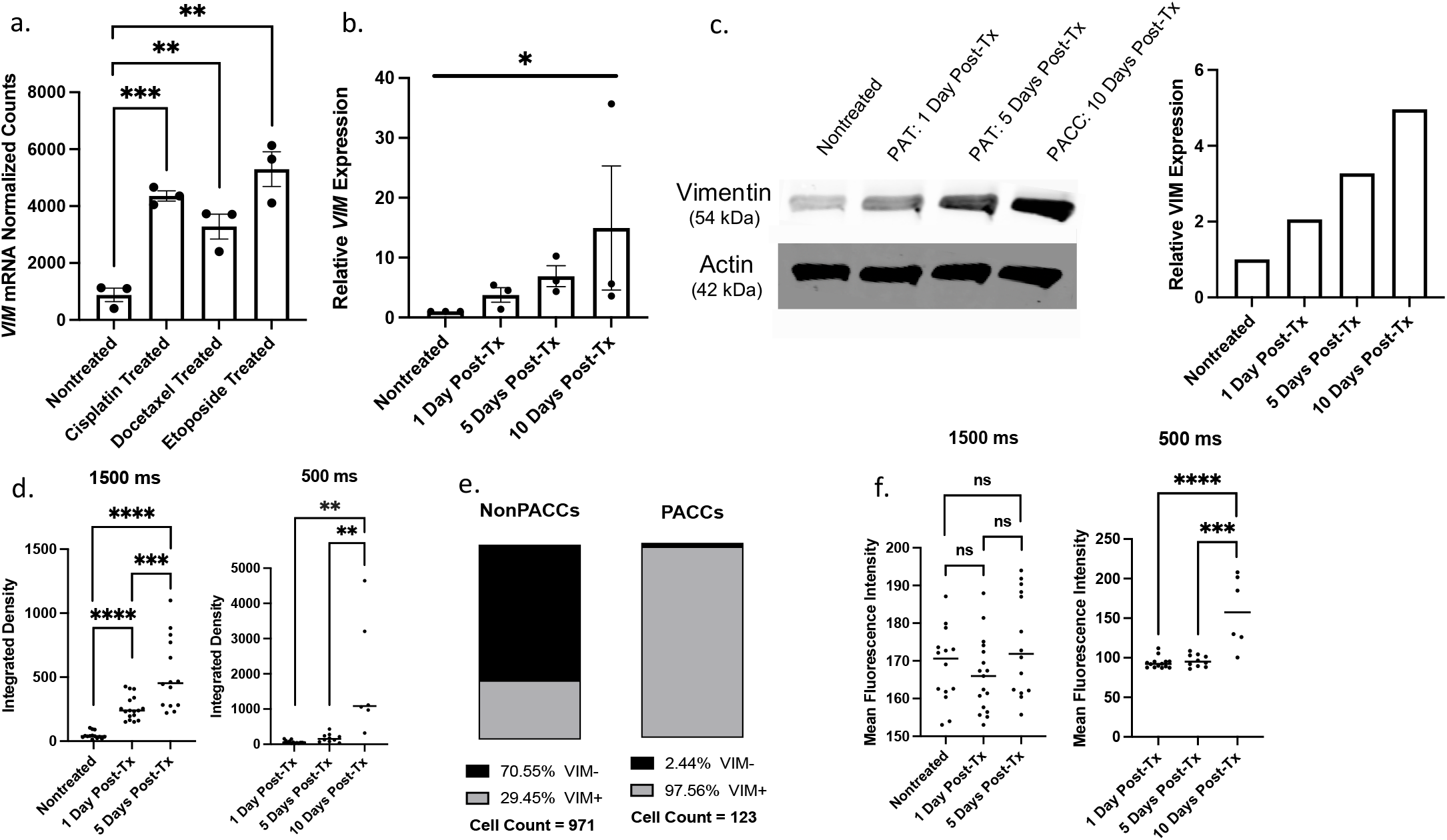
PACCs have increased expression of Vimentin at both the RNA and Protein level: a) mRNA Nanostring analysis comparing normalized counts of *VIM* mRNA transcripts in nontreated, cisplatin-treated, docetaxel-treated, and etoposide-treated populations. b) RT-qPCR analysis of relative *VIM* expression between nontreated cells, cells undergoing a PAT 1 day post-treatment, cells undergoing a PAT 5 days post-treatment, and PACC cells 10 days post-treatment. c) Western blot and quantitative densitometry of VIM expression in nontreated cells, cells undergoing a PAT 1 day post-treatment, cells undergoing a PAT 5 days post-treatment, and PACC cells 10 days post-treatment. d) Integrated density of single-cell VIM signal in nontreated cells, cells undergoing a PAT 1 day post-treatment, cells undergoing a PAT 5 days post-treatment, and PACC cells 10 days post-treatment using either a 1500 ms exposure time or a 500 ms exposure time. e) Percentage of VIM positive cells by immunofluorescence tiled imaging in nonPACC parental and PACC populations. f) Mean Fluorescent Intensity of single-cell VIM signal in nontreated cells, cells undergoing a PAT 1 day post-treatment, cells undergoing a PAT 5 days post-treatment, and PACC cells 10 days post-treatment using either a 1500 ms exposure time or a 500 ms exposure time

Immunofluorescent staining was used to assess VIM expression at the single-cell level. Total per-cell VIM expression as determined by Integrated Density (i.e. summation of fluorescence intensity in all pixels of a defined cell) was significantly higher in cisplatin-induced cells than in nontreated controls, and continued to steadily increase with time post-treatment throughout the PAT and into the definitive PACC state (10 days post-treatment) (p value = 0.0360) (Figure 5d), recapitulating the data obtained via western blot. Of note, the difference in VIM staining intensity between treated cells 10 days post-treatment (PACCs) and nontreated controls (nonPACC parental cells) was appreciably different between the two groups, limiting the visualization and imaging with the same exposure time. For example, an exposure time that is sufficient to visualize VIM+ signal in nontreated controls (1500 ms) was too long for treated cells 10 days post-treatment, resulting in an oversaturated and unanalyzable image. Accordingly, a reduction in exposure time to achieve an appropriately saturated image of the treated cells 10 days post-treatment (500 ms) renders the signal of the nontreated controls invisible (Supplemental Figure 2). Although these intensity differences between the two groups limited their direct comparability, it was apparent that treated cells undergoing the PAT and entering the final PACC showed markedly increased VIM expression.

Quantification of VIM positivity in definitive PACCs and nonPACC parental cells showed that 98% of cells in the PACC state were VIM+ (120/123) compared to 29% (286/971) of nonPACC parental cells (Figure 5e). The localization pattern of VIM within VIM+ cells also differed between PACCs and nonPACC parental cells. In nonPACC parental cells, VIM staining showed intensely bright and punctate or clustered peri-nuclear signal. In contract, VIM staining showed more dim and diffuse filamentous networks of fibers spanning the entire cytoplasm in PACCs. In addition to visual approximations, this localization difference can also be quantified by the Mean Fluorescent Intensity (MFI, or average fluorescence intensity per pixel among all pixels of a defined cell). Again, images were taken and analyzed using two different exposure times (1500 ms and 500 ms) to accurately capture both nontreated cells and treated cells 10 days post-treatment. Specifically, MFI quantifications showed no significant differences between nontreated cells and two groups of treated cells in various stages of the PAT analyzed after various number of days post-treatment (Figure 5f). Treated cells analyzed 10 days post treatment (definitive PACCs) did have an increased MFI compared to nontreated (nonPACC parental) cells, indicating that VIM overexpression dominates VIM content dynamics more so than VIM redistribution or re-localization does by 10 days post-treatment (Figure 5f). These VIM localization findings match that of other recently published work [71].

### The Vimentin content of cells in the PACC state does not correlate with their motility

To assess the role of VIM in PACC motility, we repeated the chemotactic migration experiment and then fixed and performed IF for VIM protein immediately following the final captured time lapse image. We compared end-point VIM expression and motility parameters of PACCs by linear regression. There was no correlation between measures of VIM Integrated Density (i.e. total VIM content per cell) and any motility parameter analyzed, including Euclidean distance travelled (R squared = 0.0009918), directness of PACC movement (R squared = 0.007451), or accumulated distance travelled (R squared = 0.0005488) (Figure 6a-c). Similarly, there was no correlation between VIM MFI (i.e. surrogate approximate of VIM localization) and any motility parameter analyzed. (Euclidean distance R squared = 0.01333) (Directness R squared = 0.005681) (Accumulated distance R squared = 0.04932) (Figure 6d-f).

**Fig. 6.**
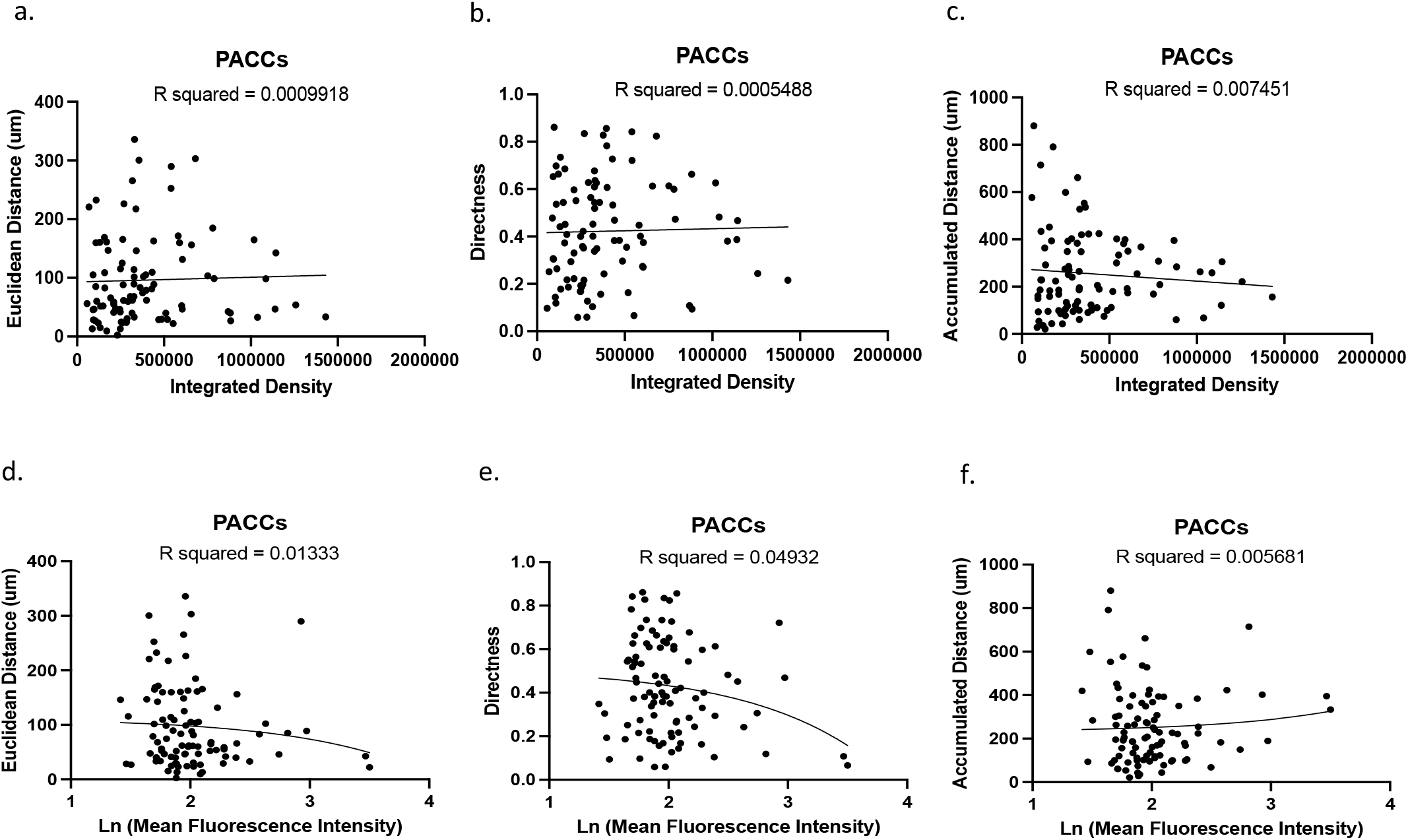
PACC Vimentin content does not correlate with PACC motility: a-c) Linear regression comparing the Integrated Density of VIM signal by immunofluorescence imaging to either the A) Euclidean distance travelled, B) directness of motility of, or C) accumulated distance travelled by PACCs. d-f) Linear regression comparing the Mean Fluorescence Intensity of VIM signal by immunofluorescence imaging to either the A) Euclidean distance travelled, directness of motility of, or C) accumulated distance travelled by PACCs

### Cells in the PACC state are equally as anoikis resistant as nonPACC parental cells

In the third step of the metastatic cascade, CTCs must survive long enough within the circulatory system to reach capillary beds supplying a distant secondary organ site. A primary threat to CTC survival in the circulation in anoikis, or programmed cell death in response to cell:ECM detachment. Anoikis resistance of PACCs and nonPACC parental cells was evaluated using a serial low-adhesion tissue culture plate to normal-adhesion tissue culture plate re-adhesion assay followed by metabolic cell viability quantification. Cells in the PACC state were equally as anoikis resistant as nonPACC parental cells (Figure 7a). Alternative quantification using DNA staining also showed equal anoikis resistance between the two groups (Data not shown).

**Fig. 7.**
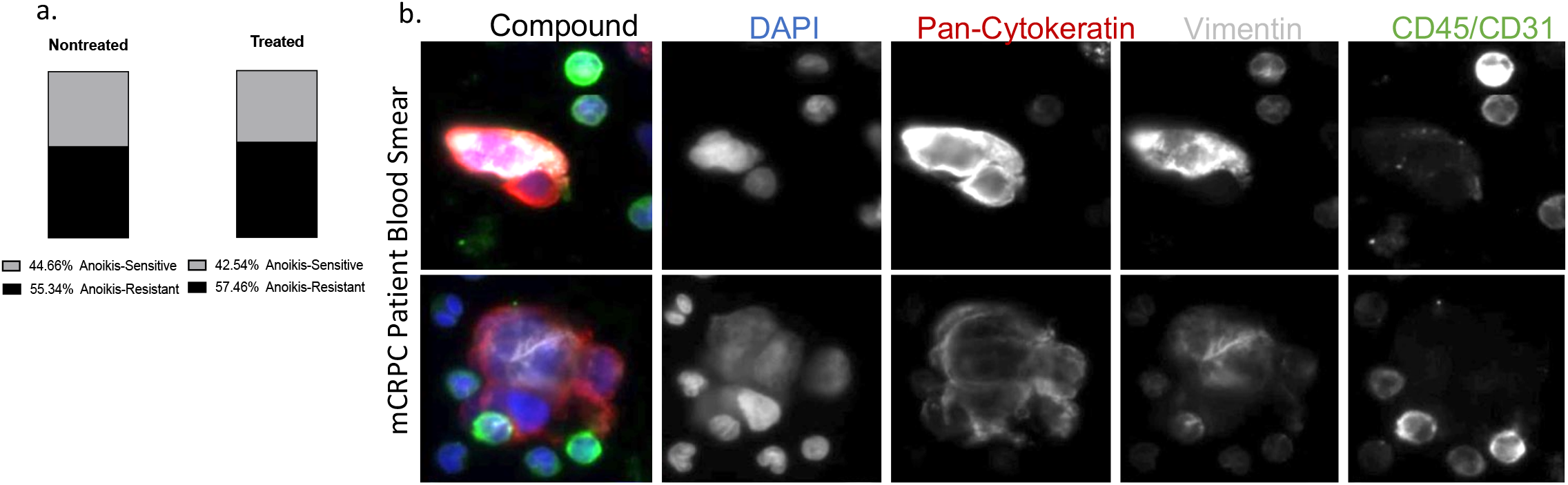
PACCs can survive in the circulatory system: a) Quantification of anoikis resistance in nontreated cells and treated cells 5 days post-treatment. Acquired using a serial low-adhesion tissue culture plate to normal-adhesion tissue culture plate re-adhesion assay. b) Images of physically enlarged blood cells collected from a patient with metastatic castrate-resistant prostate cancer (mCRPC) with increased genomic content. The cells are positive for pan- cytokeratin signal and Vimentin signal, but negative for CD45 signal, indicating that the cells are PACCs.

While cells in the PACC state have been evaluated in both primary and metastatic tumors in patient samples, presence of PACCs in circulation as CTCs has not been previously shown, likely due to the exclusion of large-nucleated cells by most automated IF-based CTC algorithms. By utilizing a selection-free CTC platform and modified analysis to include polyploid cancer cells, cells in the PACC state were identified in the circulation of a patient with metastatic castrate-resistant prostate cancer (mCRPC). This is critical data indicating presence of PACCs among CTCs *in vivo* (Figure 7b).

## Discussion

There are barriers to successful metastasis at every step of the metastatic cascade. Only metastasis-competent cells can successfully invade the primary organ, intravasate into the circulation, survive, extravasate into a distant secondary site, and then colonize there to form a clinically detectable micrometastatic lesion. Superior completion of specific steps of the metastatic cascade is less important than complete metastasis-competence. For example, supremely invasive cancer cells may never seed lethal metastatic lesions if they are incapable of surviving within the circulation. Identification of the rare subtype of completely metastasis-competent cancer cells remains one of the most important goals of cancer research.

Several features of PACCs indicate they may be metastatically competent. Previously, PACCs have been detected in patient primary tumors and metastatic lesions, and we show evidence of PACCs in the circulations as CTCs. Here, we show that PACCs demonstrate kineses-directed motility, deformability, and anoikis resistance, three factors that predict successful invasion, intravasation, survival in the circulation, and extravasation. Furthermore, the transience of the PACC state predicts successful distant site colonization following PACC depolyploidization (i.e. generation of proliferative nonPACC cancer cell progeny of normal physical size and typical genomic content). Additionally, we also present a potential role for the cytoskeletal filament Vimentin in driving metastasis relevant PACC phenotypes.

Specifically, we conclude that cells in the PACC state are more motile than nonPACC parental cells. The large and dynamic size of PACCs limits the use of traditional Boyden chamber assays and scratch-wound assays to measure motility. We used single-cell tracking following time-lapse microscopy to measure multiple movement parameters, including i) net Euclidean distance travelled and ii) directness of movement. Euclidean distance is a more metastasis-relevant measure of invasive potential than accumulated distance; it captures net distances travelled away from primary tumors (a cancer cell’s initial location) that cannot be captured by measures of accumulated distance. Other groups studying PACC motility in an adjacent cancer cell models have also shown that cells in the PACC state demonstrate increased motility and increased persistence (i.e. an alternative measure of directness) compared to nonPACC parental cells [71]. Though it is conceivable that the increased size of PACCs could cause a proportional increase in their “step size,” we find that any such scaling factor cannot explain the observed increases in PACC motility: there is absolutely no correlation between PACC cell area and PACC motility parameters. Rather, this suggests cancer cells activate specific motility programs as they undergo PAT in response to applied stress.

In addition to quantifying motility, we also performed qualitative analyses of PACC migration movies. Such analyses showed that PACCs primarily perform mesenchymal-type migration. In this movement type, the cells are flat and inch along using an alternating combination of pseudopodia elongation and trailing edge contraction. PACCs with flat morphology can also move using a combination of smooth ruffling and gliding motions without obvious pseudopod formation or extension. Occasionally, clear ameboid-type migration can also be observed in PACCs, in which rounded-up cells bleb continuously and amorphously while traveling very quickly across the field of view [72, 73].

It is useful to study cancer cell motility in the context of the habitat selection theory of (organotropic) cancer metastasis [56]. Habitat selection uses both classic cancer and ecology modeling to emphasize the collateral importance of motility and environment sensing in metastasis-competence, positing that metastasis-competent cells engage in kineses- directed movement. This theory predicts that emigration beyond the primary tumor is driven by the resource-guided movements of cancer cells in search of resources. When considered in the context of the nutrient-depleted, hypoxic, and acidic tumor microenvironment, the ability to directionally respond to chemotactic resource gradients offers a clear adaptive advantage. Expressly, it asserts that metastasis competency requires both motility and resource-sensing capabilities. Application of the habitat selection model to our motility experimental framework revealed that cells in the PACC state directionally respond to a 0% to 20% FBS chemotactic gradient. For example, PACCs in uniform serum-free conditions travelled in nonspecific, unoriented directions. Addition of an FBS gradient re-oriented the direction of travel toward the higher FBS concentration, while simultaneously preserving the PACC’s increased net Euclidean distance travelled and directness over nonPACC parental cells. This chemotactic ability appears to be gained as cells undergo PAT, as the directional movement of nonPACC parental cells is not influenced by the same gradient. Two statistical analyses of chemotaxis showed that PACCs exhibit directional chemotaxis in response to an FBS gradient but that nonPACC parental cells do not. As such, PACCs are either i) better able to sense presence of an FBS gradient, ii) better adapted to directionally respond to one, or iii) both. All the above would contribute to metastatic competency.

Interestingly, we found that PACCs in serum-free media conditions travelled the furthest net distance compared to PACCs exposed to other FBS conditions. This difference was not observed in nonPACC parental cells. Similarly, PACCs in serum-free media conditions trended toward traveling the most directly compared to PACCs in other FBS conditions, though this difference is not statistically significant. Again, this difference was not observed in nonPACC parental cells. These observations suggest that PACCs can sense and respond to environments devoid of nutrients by initiating direct movement (in any direction) away from their current nutrient-depleted location. This observation aligns with principles of optimal foraging theory, an ecological paradigm used to predict and describe how an animal behaves when searching for food. When no resources are available in an organism’s current location, it is most advantageous to pick a single direction and travel in it a long distance to increase the likelihood of encountering a new environment with adequate resources [56, 74, 75].

When a cell migrates, it experiences both tension and compression. The peripheral cytoplasmic region of a motile cell experiences tension (“pulling”) forces, particularly when employing mesenchymal-type migration. Simultaneously, the nucleus (the largest and least amorphous organelle in a cell), experiences compression (“pushing”) forces. Functional deformability is important during both invasion and intravasation/extravasation when motile cells experience both tension and compression as they maneuver through densely packed tumor cells, extra-cellular matrices, and endothelial cells [65]. Notably, nuclear integrity is the most important determinant of cellular viability in a cell experiencing deformation.

It is useful to study cancer cell deformability and nuclear integrity in the context of molecular biophysics, particularly using stress-strain diagrams. Stress-strain diagrams depict the causal relationship between applied forces (the stress) and resultant shape deformation (the strain) of an object. A hyper-elastic biomaterial is one with a nonlinear stress-strain relationship that undergoes extremely large deformations in response to minimal amounts of applied force. Hyper- elastic biomaterials are nearly incompressible, meaning they can change their shape while retaining near-constant volume. Furthermore, they readily return to their original shape when the force is removed. Most notably, hyper-elastic biomaterials stiffen dramatically when under intense compression, but exhibit softening when under tension [76].

Though the MTC and AFM data initially appear to contradict one another, a molecular biophysical analysis reveals that together, they indicate that PACCs are hyper-elastic. In the MTC assay, a magnetic field exhibited a peripheral pulling force on the cytoskeleton, creating tension. During MTC, PACCs demonstrated decreased intracellular cytoskeletal network stiffness compared to nonPACC parental cells, which aligns with the characteristics of a hyper-elastic cytoskeleton experiencing tension. In the AFM assay, a pointed probe was depressed into the perinuclear region with uniform downward force, creating compression. During AFM, PACCs appeared to be, on average, equally as cortically stiff in the perinuclear region as nonPACC parental cells, though there did exist some cells in the PACC state with extremely stiff peri-nuclear regions far exceeding the stiffness of any nonPACC parental cell. The PACCs’ maintenance of (or occasional increase in) cortical stiffness aligns with the characteristics of a hyper-elastic cytoskeleton experiencing compression. The hyper-elastic nature of PACCs provides them with both i) a greater capacity for metastasis-promoting peripheral deformability than nonPACC parental cells and ii) equal or greater nuclear integrity than nonPACC parental cells. These two conclusions thus support a model of PACCs as metastasis-competent cells.

Previous published work shows that VIM intermediate filaments are hyper-elastic and thus allow cells, and particularly nuclei, to withstand extreme deformations without fracture or rupture [77]. Overall, this simultaneously confers both intracellular and cortical cytoskeletal strength and stretchability. All PACCs show increased *VIM*/VIM content at the RNA and protein level when compared to nonPACC parental cells by four orthogonal techniques.

Recent work published by Dawson et. al also showed that VIM content was increased in PACCs in a breast cancer MDAMB231 paclitaxel-induced PACC model [71, 78, 79]. Additionally, they showed that increased VIM content in their model is necessary for migratory persistence, or enhanced directness of PACC motility. Chemical inhibition (with low-dose acrylamide) or siRNA-mediated knockdown of *VIM* resulted in both decreased PGCC spreading/surface area and decreased migratory persistence. This work suggested that in their models, VIM played a key role in orienting the polarization of a PGCC performing mesenchymal-type migration. Contrarily, our observed lack of correlation between VIM abundance and PACC motility suggests that the increased VIM content in our PACCs primarily plays a role in supporting deformability and cytoskeletal toughness via its hyper-elastic properties.

Following invasion and intravasation, a metastatically competent cell must survive its time in the circulation. Deformability is pertinent for survival in the circulation. CTCs must resist the shear stress of blood flow to survive the circulatory system. Increased deformability heightens a cell’s ability to withstand shear stresses of blood flow, as has been observed in studies of red blood cell dynamics [80]. CTCs must also resist anoikis. Our anoikis resistance assays show that PACCs are equally as anoikis resistant as nonPACC parental cells. Considering that the PC3 cell line was derived from an osseous metastatic lesion of a prostate cancer, it is unsurprising that PC3 nonPACC parental cells boast a basal high level of anoikis resistance. We’ve observed that PC3 cells maintain this level of anoikis resistance as they undergo PAT to enter the PACC state, supporting the model of PACCs as metastasis-competent cells. Identification of PACC CTCs within the blood of a patient with metastatic castrate-resistant prostate cancer also supports this model.

The observed motility, environment-sensing, and deformability in PACCs also predict competent extravasation by PACCs. The habitat selection model highlights the requirements of a CTC halted in the capillary of a distance secondary organ to i) sense presence of an appropriate resource within the tissue of the secondary organ and ii) directionally respond by travelling out of the circulation and into the secondary organ [56], during which it must deform as it moves between endothelial cells. Thus, the properties that predict a PACC’s invasion and intravasation competency are the same ones that predict a PACC’s extravasation competency. Lastly, the PACC state is an inherently transient one. PACCs undergo eventual depolyploidization to produce proliferative progeny of typical cancer cell and genomic content size. As such, PACCs retain the capability of distant second site colonization (the fifth and final step required for metastatic competency).

In conclusion, identification of the rare subpopulation of metastasis competent cells remains critically important. Several clinical and experimental observations suggest PACCs may be metastatically competent. In this work, we show that PACCs demonstrate metastatic-competent motility, environment-sensing, deformability, and anoikis-resistance. These characteristics, alongside the PACCs inherent progeny-forming potential, support the model of PACCs as metastasis-competent cells, and in the least indicate they are novel candidates for continued *in vivo* investigation.

## Methods and Materials

### Cell Culture

Cell culture experiments were performed with the PC3-luc prostate cancer cell line [81]. All cells were cultured with RPMI 1640 media with L-glutamine and phenol red additives. (Gibco) supplemented with 10% Premium Grade Fetal Bovine Serum (Avantor Seradigm) and 1% 5000 U/mL Penicillin-Streptomycin antibiotic (Gibco), at 37 degrees Celsius and in 5% CO2. PC3 cells were authenticated and tested for *mycoplasma* biannually (Genetica).TryplE Express Enzyme without phenol red additive was routinely used as a dissociation reagent unless otherwise stated (Gibco), and all centrifugations were performed at 1000 xg for 5 minutes unless otherwise stated.

### PACC induction

Cells were plated at a density of 625,000 cells per T75 flask. 24 hours after plating, cells were treated with 6 uM (GI50) of the chemotherapeutic drug cisplatin (resuspended in PBS with 140mM NaCl) for 72 hours, unless otherwise indicated (Millipore Sigma). After 72 hours of treatment, drug-containing media was removed and replaced with fresh media. Treated cells were kept in culture for up to an additional 10 days, at which point they are considered definitive PACCs. Fresh media was replenished every 4 days. Images of cells undergoing a PAT following the PACC-induction protocol were taken from day 0 through day 15 using the Incuctye S3 Live Cell Analysis System (Sartoris) with daily media changes following removal of treatment.

To ensure purity of the PACC population for bulk-cell experiments (Western Blot, RT-qPCR), any nonPACC cells remaining in culture following chemotherapeutic PAT induction are excluded via size-based filtration. Treated cells are dissociated from the flask, resuspended in 25 mLs media, and flowed through 15 micron filters (PluriSelect) using back- pressure from an attached 5 mL luer-lock syringe (BD) (5mL cell suspension/per filter). Cells caught in the filters are then collected in 10 mL media. Collected cells are spun down at 1000 xg for 5 minutes and can be reused/replated in any capacity.

### Flow Cytometry

Flow cytometry using FxCycle Violet stain (Invitrogen) for analysis of DNA content was performed on 1,000,000 non treated cells and 1,000,000 treated cells immediately following release from chemotherapeutic treatment. The stain was applied according to the manufacturer’s protocol and analyzed using the Attune NxT Flow Cytometer (Thermo Fisher). Live-cell and doublet-exclusion gating was performed unstained controls. Analysis was performed using FlowJo.

### Immunofluorescence

Treated or nontreated cells were plated on glass-bottom chamber slides (Falcon, Corning) at various days (1, 5, or 10 days) following release of chemotherapeutic treatment and allowed to adhere overnight. Cells were then fixed with cold 4% methanol-free PFA (Thermo Scientific), for 15 minutes at room temperature and then washed 3 times for 5 minutes each with pH 7.4 Phosphate Buffered Saline (PBS) (Gibco). Cells were then simultaneously permeabilized and blocked with a solution of 0.25% Surfact-Amps X-100 (Thermo Scientific) in PBS and 5% Normal Goat Serum (Abcam) in PBS for 60 minutes at room temperature. Cells were then incubated in primary antibody Vimentin (D21H3) XP Rabbit mAb (Cell Signaling Technologies) diluted 1:200 OR Beta-Actin (8H10D10) Mouse mAb (Cell Signaling Technologies diluted 1:5,000 in a solution of 1% Bovine Serum Albumin (BSA) Fraction V (Fisher Scientific) and 0.25% Surfact-Amps X-100 in PBS overnight at 4 degrees Celsius. Cells were washed 3 times for 5 minutes each with PBS and incubated in secondary antibody Goat anti Rabbit IgG H+L Cross-Absorbed Secondary Antibody, Alexa Fluor 488 (Invitrogen) OR Goat anti Mouse IgG H+L Cross-Absorbed Secondary Antibody, Alexa Fluor 555 (Invitrogen) diluted 1:200 in a solution of 1% BSA and 0.25% Surfact-Amps X-100 in PBS for 3 hours at room temperature. Cells were then washed 3 times for 5 minutes each with PBS and mounted using ProLong Diamond anti-fade mountant with DAPI (Invitrogen) and allowed to cure overnight. All slides were imaged using an AxioCam Mrm (Zeiss) camera on an Observer.Z1 microscope (Zeiss) using the XCite 120Q fluorescence illuminator (Excelitas) and analyzed using ImageJ image analysis software.

### Migration Assays

Chemotactic gradients were established using the µ-Slide Chemotaxis system (Ibidi) following the manufacturer’s protocols. NonPACC parental cells were seeded at 3,000,000 cells per mL of media. PACCs were seeded at 500,000 cells per mL. Positive control cells were seeded in 10% FBS-containing media, were allowed to adhere overnight, were rinsed twice with PBS before uniform solution of 20% FBS-containing media was applied across the µ-Slide. Negative control cells were seeded in serum-free media, were allowed to adhere overnight, were rinsed twice with PBS before uniform solution of serum-free media was applied across the µ-Slide. For the gradient experiment, cells were seeded in serum-free media, were allowed to adhere overnight, were rinsed twice with PBS before a 0-20% chemotactic gradient of FBS-containing media was applied across the µ-Slide. Immediately after addition of the gradient, the cells were imaged via live-cell time lapse microscopy using the EVOS FL Auto Imaging System (Life Technologies). Images were taken with a 10X objective every 30 minutes for 24 hours. Environment chamber conditions were 37 degrees Celsius, 5% CO2, and 20% O2. Immediately following time lapse microscopy, cells were processed for Immunofluorescent labeling of Vimentin (see above).

Time lapse images were analyzed using the Manual Tracking and Chemotaxis and Migration macros in ImageJ image analysis software. All cells analyzed were randomly selected. Cells that underwent division, apoptosis, or moved out of frame were excluded from analysis. 2D surface area measurements of all cells analyzed were also obtained via ImageJ.

### Spontaneous and Forced Bead Motions with Magnetic Twisting Cytometry

NonPACC parental cells and PACCs were seeded in 96-well stripwell microplates (Corning). NonPACC parental cells were plated at 30,000 cells/well and PACCs were plated at plated at 3,000 cells/well to account for differences in cell size. RGD-coated ferrimagnetic microbeads (∼4.5□μm in diameter) were added, anchoring to the cytoskeleton via cell surface integrin receptors of adherent living cells. Spontaneous nanoscale displacement of individual microbeads (∼100 beads per field of view) was recorded at a frequency of 12 frames/s for t_max_ ∼300s via a CCD camera (Orca II-ER, Hamamatsu). Bead trajectories in two dimensions were then characterized by computing mean squared displacement of all beads as a function of time [MSD(t)] (nm_^2^_) as previously described [82]. We then applied forced motions of the functionalized microbeads using MTC as previously described [83] to measure the stiffness of individual cells. The RGD-coated ferrimagnetic microbeads were magnetized parallel to the cell plating (1,000 Gauss pulse) and twisted in a vertically aligned homogenous magnetic field (20 Gauss) that was varying sinusoidally in time. The sinusoidal twisting magnetic field caused both a rotation and pivoting displacement of the beads that lead to the development of internal stresses that resist its motion. Lateral bead displacement was optically detected with a spatial resolution of 5nm, and the ratio of specific torque to beads displacement was computed and expressed as the cell stiffness (Pa/nm). The same population of cells (with attached RGD beads) was used to acquire both the cytoskeletal rearrangement and stiffness measurements in the same experiment.

### Atomic Force Microscopy

NonPACC parental cells and PACCs were plated in 60 mm dishes. NonPACC parental cells were seeded at 80,000 cells/dish and PACCs were seeded at 10,000 cells/dish achieve roughly 25% confluency. Cells were incubated overnight to adhere. After 24 hours, AFM experiments were done with a cantilever with pointed tips made of Silicon Nitride (SiN). The nominal spring constant of the cantilever was 0.01 N/m (Bruker) on an MFP3D (Asylum Research) instrument. The Cantilever tips were calibrated every time before experiments using thermal fluctuation method [84]. To measure the apical stiffness, cells were indented using contact mode with a maximum peak force of 1 nanoNewton, to get force-displacement curves. All data processing was done using Igor pro software (Wavemetrics). The young’s modulus was obtained by fitting the force-displacement curves with Hertz model, which related the applied force (F) by the cantilever tip to the indentation (δ) and the Young’s modulus (E) using the equation below, where α is the tip opening angle (35°) and v is the Poisson ratio (which is assumed to be 0.5 for soft biological materials) [85].

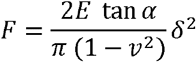

### Western Blot

Treated cells were filtered at various days (1, 5, or 10 days) following release of chemotherapeutic treatment. Each population of filtered cells as well as nonfiltered nontreated cells were pelleted and then lysed with an appropriate amount of RIPA Lysis and Extraction Buffer (Thermo Scientific) with Halt Protease and Phosphatase Inhibitor Cocktail (Thermo Scientific) for 30 mins, rotating in 4 degrees Celsius. Lysates were spun at 21,000 xg for 15 mins in 4 degrees and supernatant was stored at -80 degrees Celsius. 50 ng of protein (measured by Pierce BCA Protein Assay, following manufacturer’s protocol) (Thermo Scientific) was added to a 1:4 mixture of Laemmli Sample Buffer (BioRad) and 2- Mercaptoethanol (BioRad) and ran through a 4-20% Mini-ProTEAN TGX gel (BioRad). The gel was transferred via Trans-Blot SD Semi-Dry Transfer Cell (BioRad) onto a 0.2 micron Nitrocellulose Trans-Blot Turbo Transfer Pack using the 7 min protocol designed for Mixed Molecular Weights. The blot was blocked in Casein Blocking Buffer (Sigma-Aldrich) for 1 hour at room temperature with shaking, and then transferred to primary antibody Vimentin (D21H3) XP Rabbit mAb (Cell Signaling Technologies) diluted 1:1000 in casein and incubated overnight at 4 degrees Celsius with shaking OR Monoclonal Anti-Beta-Actin mouse antibody (Sigma) diluted 1:5000 in casein and incubated at room temperature for 1 hour. The blot was then washed 3 times for 5 minutes each with pH 7.4 Tris-Buffered Saline (Quality Biological) with 0.1% Tween 20 (Sigma) (TBST) and incubated in secondary antibody IRDye 700CW Goat anti-Rabbit (Li-Cor) OR IRDye 680RD Goat anti-Mouse (Li-Cor) diluted 1:20,000 in Casein for 1 hour at room temperature. The blot was then washed 3 times for 5 minutes with TBST and imaged using the Odyssey Western Blot Imager (Li-Cor). Densitometry analysis of images was performed using ImageJ image analysis software.

### RT-qPCR

Treated cells were filtered at various days (1, 5, or 10 days) following release of chemotherapeutic treatment. Each population of filtered cells as well as nonfiltered nontreated cells were lysed using a QIAshredder Kit (Qiagen) following the manufacturer’s protocol. RNA was extracted from lysates using an RNeasy Mini Kit (Qiagen) following the manufacturer’s protocol. RNA was converted to cDNA (1 ug RNA per reaction) using the iScript cDNA Synthesis Kit (Bio-Rad) following the manufacturer’s protocol. RT-qPCR reactions were performed using SsoFast EvaGreen Supermix (Bio-Rad) following the manufacturer’s protocols on the CFX96 Real-Time PCR Detection System (Bio-Rad). Beta-Actin was used as the housekeeping control gene. Gene expression was normalized to a housekeeping gene and calculated with the delta-delta Ct method. The following primers were used:

Vimentin Forward: 5’ TGCCGTTGAAGCTGCTAACTA 3’
Vimentin Reverse: 5’ CCAGAGGGAGTGAATCCAGATTA 3’
Actin Forward: 5’ ACGTGGACATCCGCAAAGAC 3’
Actin Reverse: 5’ CAAGAAAGGGTGTAACGCAACTA 3’

### mRNA Nanostring

Cells were treated with either 6 uM cisplatin, 5 nM Docetaxel, or 25 nM Etoposide. Treated cells were filtered 1 day following release of chemotherapeutic treatment. Each population of filtered cells as well as nonfiltered nontreated cells were lysed using a QIAshredder Kit (Qiagen) following the manufacturer’s protocol. RNA was extracted from lysates using an RNeasy Mini Kit (Qiagen) following the manufacturer’s protocol. RNA was converted to cDNA using the iScript cDNA Synthesis Kit (Bio-Rad) following the manufacturer’s protocol. The complete product was used as input for hybridization with 770 nCounter PanCancer Progression probes for 16 hours according to manufacturer’s protocols. Loaded cartridges were run on an nCounter Sprint (NanoString Technologies). Gene expression data quality control was analyzed using nSolver Analysis Software 4.0.70 (NanoString Technologies). All samples were normalized to the total counts of the nCounter- defined positive controls to reduce lane-to-lane variation from cartridge loading and normalize binding affinity across all samples surveyed. mRNA transcript reads of less than 40 were considered undetected.

### Invasion Chambers

The Polydimethylsiloxane (PDMS)-based microfluidic device was fabricated as previously described [86, 87]. The PDMS-based microfluidic devices contained a series of parallel microchannels with varying widths of 3, 6, 10, 20, and 50- µm, lengths of 200 µm, and heights of 10 µm. The microchannels were perpendicular to a 2D cell seeding area and were coated with 20 µg/ml of collagen type I at 37 °C for 60 to 80 minutes. NonPACC Parentals or PACCs were dissociated with PBS-Based Enzyme Free Cell Dissociation Buffer (Gibco). 1,000,000 cells per mL of media were loaded into the microfluidic device. Cells were imaged via live-cell time lapse microscopy using the EVOS FL Auto Imaging System (Life Technologies). Images were taken with a 10X objective every 30 minutes for 24 hours. Environment chamber conditions were 37 degrees Celsius, 5% CO2, and 20% O2. 4

### Anoikis

25,000 treated and filtered cells or 25,000 nontreated cells were simultaneously plated in i) a 12-well low-adhesion tissue culture plate (Corning) and ii) a 12-well normal-adhesion positive control tissue culture plate. After 72 hours, both the treated and nontreated cells initially plated in the low-adhesion plates were independently transferred to fresh normal adhesion plates, in which they were cultured for an additional 48 hours. Both the treated and nontreated cells initially plated in normal-adhesion plates were cultured typically for a total of 120 hours. 120 hours after initial seeding, the cellular viability was measured as a proxy for cellular number or density using the alamarBlue Cell Viability Agent (Invitrogen) according to the manufacturer’s protocols. A 2-hour incubation was used and fluorescence (excitation 560, emission 590O was measured via FLUOstar Omega plate reader (BMG Labtech). Anoikis resistance for each condition was calculated by creating a ratio of the viability of the low-adhesion plate challenged cells to the positive control normal-adhesion plated cells for cells from each treatment condition. Alternatively, 120 hours after initial seeding, the cellular density was measured using a crystal violet DNA stain, in which cells were fixed with 4% PFA for 15 minutes at room temperature, stained with 0.05% crystal violet suspended in 20% methanol for 20 minutes, and thoroughly washed with PBS. After drying, 10% Acetic Acid was used to resuspend the stain. Absorbance (596 nm) was measured using a FLUOstar Omega plate read (BMG Labtech). Anoikis resistance for each condition was calculated by creating a ratio of the absorbance of the low-adhesion plate challenged cells to the positive control normal-adhesion plated cells for cells from each treatment condition.

### CTC detection

Blood was collected from a patient diagnosed with de novo metastatic prostate cancer following castration resistance as previously described in [88]. Blood cells were plated on cell-adhesive (Marienfeld) slides and underwent immunofluorescent stained for a mixture anti-human cytokeratins 1,4,5,6,8,10,13,18 and 19, CD45, Vimentin, and DAPI as previously described in [88]. Slides were imaged and analyzed as previously described in [88].

### Statistics

Prism9 was used to generate all graphics. Nonparametric T-Tests (Mann-Whitney) were performed using Prism9 to generate all reported P values unless otherwise stated. Elsewhere, nonparametric one-way ANOVA (Kruskal Wallace) was performed using Prism9 to analyze RT-qpCR-generated data, and Rayleigh’s Tests was performed using ImageJ to analyze chemotaxis data. Throughout, an alpha value of 0.05 was used.

NS = nonsignificant = P > 0.05.

* = P < 0.05

** = P < 0.01

*** = P < 0.001

**** = P < 0.0001

## Acknowledgements

The authors thank Dr. Peter Kuhn and Dr. Jim Hicks for contributing PACC CTC detection data. They also thank members of the Pienta Amend Lab and the Brady Urological Institute for thoughtful conversation and invaluable feedback. This work was supported by the William and Carolyn Stutt Research Fund, Ronald Rose, MC Dean, Inc., William and Marjorie Springer, Mary and Dave Stevens, Louis Dorfman, and the Jones Family Foundation.

## Statements and Declarations

### Author Contributions

Conceptualization: [MMM, KJP, SRA]; Methodology: [MMM, NK, MIC, SJL]; Formal analysis and investigation: [MMM, NK, MIC, SJL]; Writing - original draft preparation: [MMM]; Writing - review and editing: [MMM, SJL, SAS, KJP, SRA]; Funding acquisition: [SAS, SXS, KK, KJP, SRA]

## Supplemental Information

**Online Resource 1**

Video of nonPACC parental cell motility created from 10X timelapse images taken at 30-minute intervals for 24 hours.

**Online Resource 2**

Video of PACC motility created from 10X timelapse images taken at 30-minute intervals for 24 hours.

**Online Resource 3**

Video of PACC functional deformability through a 10 um invasion channel, in response to a 0-20% FBS gradient created from 10X timelapse images taken at 30-minute intervals for 48 hours.

**Online Resource 4**

Video of PACC functional deformability through a 20 um invasion channel, in response to a 0-20% FBS gradient created from 10X timelapse images taken at 30-minute intervals for 48 hours.

