## Supplemental Figures for "Cells in the Polyaneuploid Cancer Cell (PACC) state have increased metastatic potential"

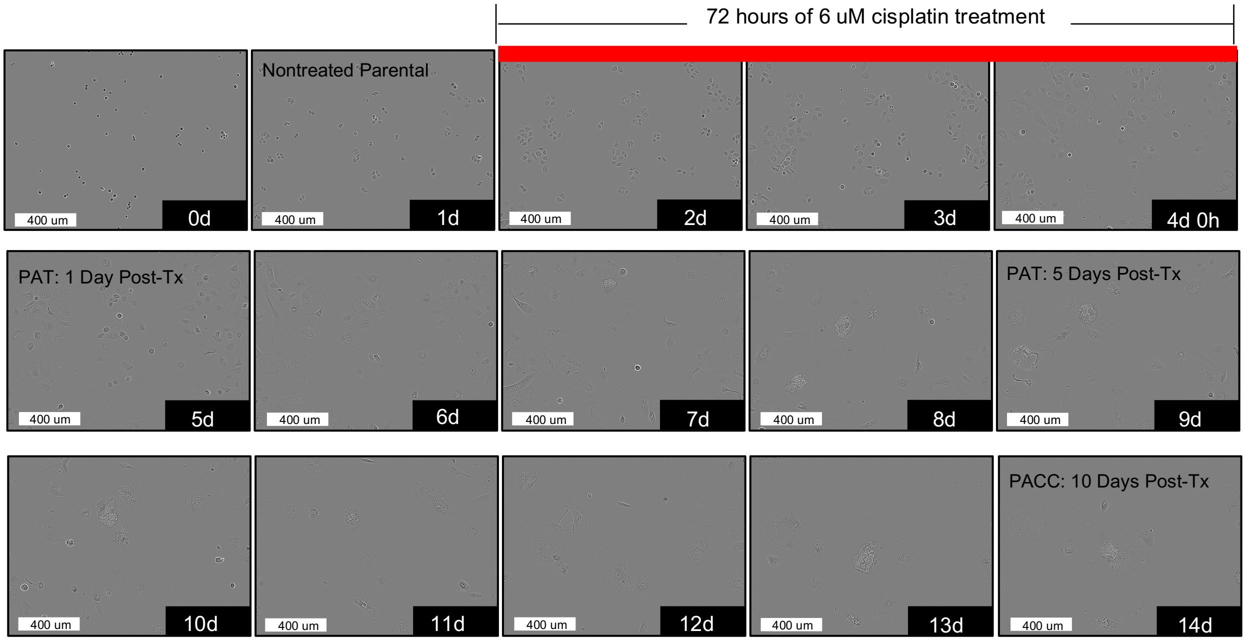


**Supplemental Fig. 1**
One representative frame of PC3 cells subjected to cisplatin-medicated PACC generation throughout a 14 day, 10X in-incubator time lapse.

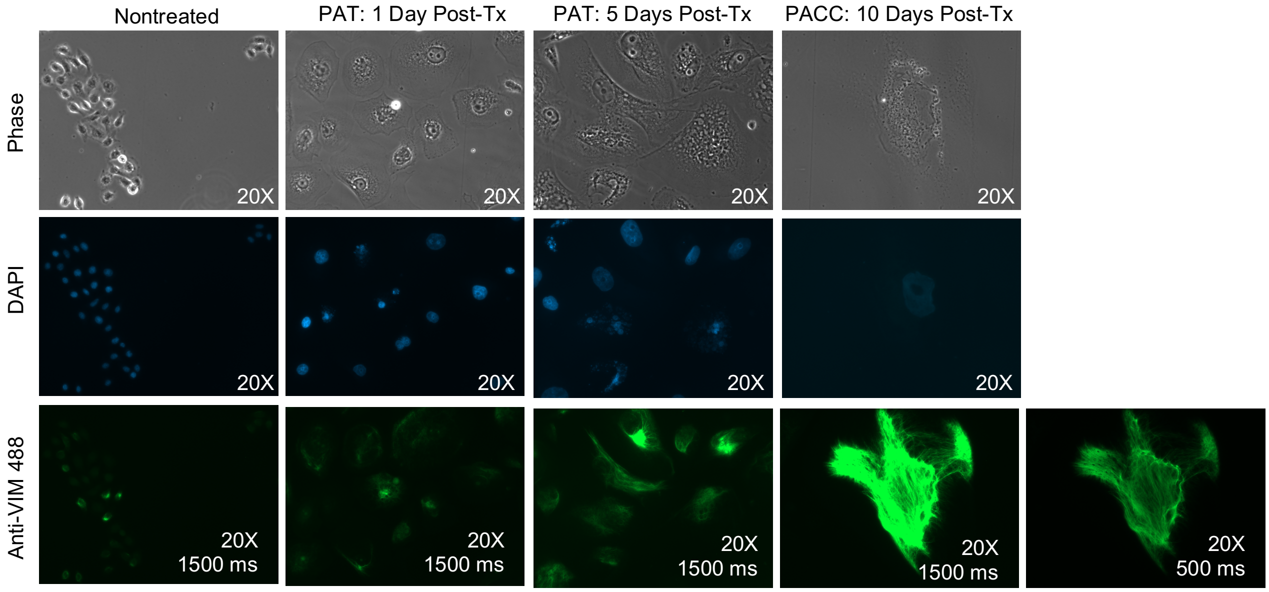


**Supplemental Fig. 2**
Representative 20X immunofluorescent images of nontreated cells, cells undergoing a PAT 1 day post-treatment, cells undergoing a PAT 5 days post-treatment, and PACC cells 10 days post-treatment co-stained with DAPI and Anti-VIM 488. Anti-VIM 488 images were acquired at two exposure times to optimize signal intensity across cell populations.
